# High content Image Analysis to study phenotypic heterogeneity in endothelial cell monolayers

**DOI:** 10.1101/2020.11.17.362277

**Authors:** Francois Chesnais, Jonas Hue, Errin Roy, Marco Branco, Ruby Stokes, Aize Pellon, Juliette Le Caillec, Eyad Elbahtety, Matteo Battilocchi, Davide Danovi, Lorenzo Veschini

**Affiliations:** Academic centre of reconstructive science, Faculty of Dentistry Oral & Craniofacial Sciences, King’s College London, Guy’s Hospital, Great Maze Pond, London SE1 9RT, UK; Centre for Stem Cells and Regenerative Medicine, King’s College London, Guy’s Hospital, Floor 28, Tower Wing, Great Maze Pond, London SE1 9RT, UK; Centre for host-microbiome interactions, Faculty of Dentistry Oral & Craniofacial Sciences, King’s College London, Guy’s Hospital, Great Maze Pond, London SE1 9RT, UK; bit.bio, Babraham Research Campus, The Dorothy Hodgkin Building, Cambridge CB22 3FH, UK

## Abstract

Endothelial cells (EC) are heterogeneous across and within tissues, reflecting distinct, specialised functions. EC heterogeneity has been proposed to underpin EC plasticity independently from vessel microenvironments. However, heterogeneity driven by contact-dependent or short-range cell-cell crosstalk cannot be evaluated with single cell transcriptomic approaches as spatial and contextual information is lost. Nonetheless, quantification of EC heterogeneity and understanding of its molecular drivers is key to developing novel therapeutics for cancer, cardiovascular diseases and for revascularisation in regenerative medicine.

Here, we developed an EC profiling tool (ECPT) to examine individual cells within intact monolayers. We used ECPT to characterise different phenotypes in arterial, venous and microvascular EC populations. In line with other studies, we measured heterogeneity in terms of cell cycle, proliferation, and junction organisation. ECPT uncovered a previously under-appreciated single-cell heterogeneity in NOTCH activation. We correlated cell proliferation with different NOTCH activation states at the single cell and population levels. The positional and relational information extracted with our novel approach is key to elucidating the molecular mechanisms underpinning EC heterogeneity.

**Summary statement:** Endothelial cells heterogeneity is key to complex collective functions and cell behaviour. We developed a novel image based endothelial cell profiling tool and quantified heterogeneity in NOTCH signalling in monolayers.

## Introduction

Endothelial cells (EC) form the inner layer of blood and lymphatic vessels and play major roles in tissue development, angiogenesis, inflammation, and immune cell trafficking (Potente et al., 2011). EC are functionally plastic and rapidly adapt to changes in the environment to preserve homeostasis. Local EC dysregulation is a hallmark of diseases such as atherosclerosis, ischemia and cancer (Park-Windhol and D’Amore, 2016). EC from different tissues and vascular beds (e.g., arteries, capillaries, veins) exhibit distinct metabolism, morphology, and gene expression (Augustin and Koh, 2017) and contribute in diverse ways to tissue development and regeneration (Itkin et al., 2016; Kusumbe et al., 2014). It is well established that EC are phenotypically heterogenous not only among different tissues, reflecting specialised organ-specific functions (Rafii et al., 2016), but also within the same tissue. Maintenance of endothelial homeostasis depends on new EC substituting senescent cells and the role of endothelial progenitor cells with high repopulating potential has been highlighted in large vessels endothelia (Yoder, 2018).

Inter-endothelial adherens junctions (IEJ) are dynamically regulated by VE cadherin (CDH5) shuffling between the cell membrane and intracellular compartments. This process presents variations across vascular beds (Augustin and Koh, 2017) and involves molecular mechanisms including VEGF receptors, cytoskeletal proteins and NOTCH family members. VEC and NOTCH signalling are well studied in angiogenesis and development. Nonetheless, the role of these molecular mechanisms in EC monolayer maintenance is less clear and this knowledge is essential to understand vessel homeostasis in different organs in the human body. So far, a comprehensive analysis of EC cultures exploring and quantifying phenotypic variance has proven prohibitively difficult because of lack of adequate tools.

Single cell phenotyping has identified and characterised intermediate cell states (MacLean et al., 2018; Siu et al., 2020) and demonstrated correspondence between phenotypes and function (Dueck et al., 2016). However, challenges in discriminating functional phenotypic variance from biological noise have emerged (Eling et al., 2019). Single cell transcriptomic (sc-OMICS) data is becoming available (mostly in mouse) (Kalucka et al., 2020) and could in principle advance the characterisation of human EC (Tikhonova et al., 2019) and provide an overview of molecular processes in distinct EC populations. Nonetheless, sc-OMICS data lacks spatial information which is essential to map variable phenotypes to function and is also required to understand higher-level functions depending on cell-cell connectivity. Of note, EC cell cycle and rapid adaptative phenomena, such as rapid increase of endothelial permeability upon VEGF stimulation are regulated by built-in sensing mechanisms that depend on cell-cell interaction (Acar et al., 2008). Importantly, EC monolayers maintain their integrity over years while exerting a variety of system-level functions which are emerging properties of cells in contact (McCarron et al., 2017). It has been proposed that endothelial adaptability and diversity of functions within an EC population depends on cell heterogeneity (McCarron et al., 2019).

To examine individual cell heterogeneity and extract spatial information from EC monolayers, we developed an endothelial cell profiling tool (ECPT) based on high-content image analysis (HCA). ECPT captures a wealth of cellular, subcellular, and contextual information enabling extensive characterisation of cell cycle and IEJ. This unbiased approach allows quantification of EC heterogeneity measuring feature variance (Eling et al., 2019). Taking advantage of machine learning based methods, we performed automated and accurate classification of IEJ using junctional CDH5 immunostaining, and evaluation of NOTCH activation at the single cell level. In total, we analysed data from >20000 images across 9 independent experiments detailing selected measurements from >300,000 cells including individual junction objects. Overall, we present 1) a novel tool for single EC profiling at a previously inaccessible scale, 2) the validity of this analysis to quantify previous observations and 3) new key relationships across features that are distinct between different EC types.

## Results

### Challenging quantification of endothelial cell heterogeneity by high-content imaging

While heterogeneity of EC has been proposed, few transcriptomic studies have successfully quantified the phenotypic variance of EC within the same vascular bed (Chavkin and Hirschi, 2020). We previously reported the value of specific Vascular Endothelial Cadherin (CDH5) and NOTCH quantification to benchmark EC in intact monolayers (Wiseman et al., 2019) and now extend our observations to include relational information between individual cells. These cannot currently be retrieved with single cell transcriptomic analysis. We cultured primary HUVEC (Human Umbilical Vein EC), HAoEC (Human Aortic EC) and HPMEC (Human Pulmonary Microvascular EC) for 48 or 96 hours in the absence or presence of VEGF. We aimed for sufficient cells to reach confluency upon adhesion and spreading (~24 h) and we cultured for a further 24 or 72 hours to enable formation of stable IEJ. EC demonstrate a uniform cobblestone morphology under low magnification, confirming monolayer stabilisation.

Immunostaining and live imaging of EC monolayers has highlighted differences in cell morphology and junction patterns in response to stimuli or under distinct cell culture conditions (Abu Taha et al., 2014; Lampugnani et al., 1992; Seebach et al., 2016; Vestweber et al., 2009). We focussed on analysing proliferation, junctions, and NOTCH activation. We stained for CDH5, NOTCH1 (or actin cytoskeleton), HES1 (a NOTCH target gene) and nuclei. In line with previous literature, qualitative inspection at high magnification revealed remarkable morphological differences between the cell types in culture (Figure 1).

**Figure 1:**
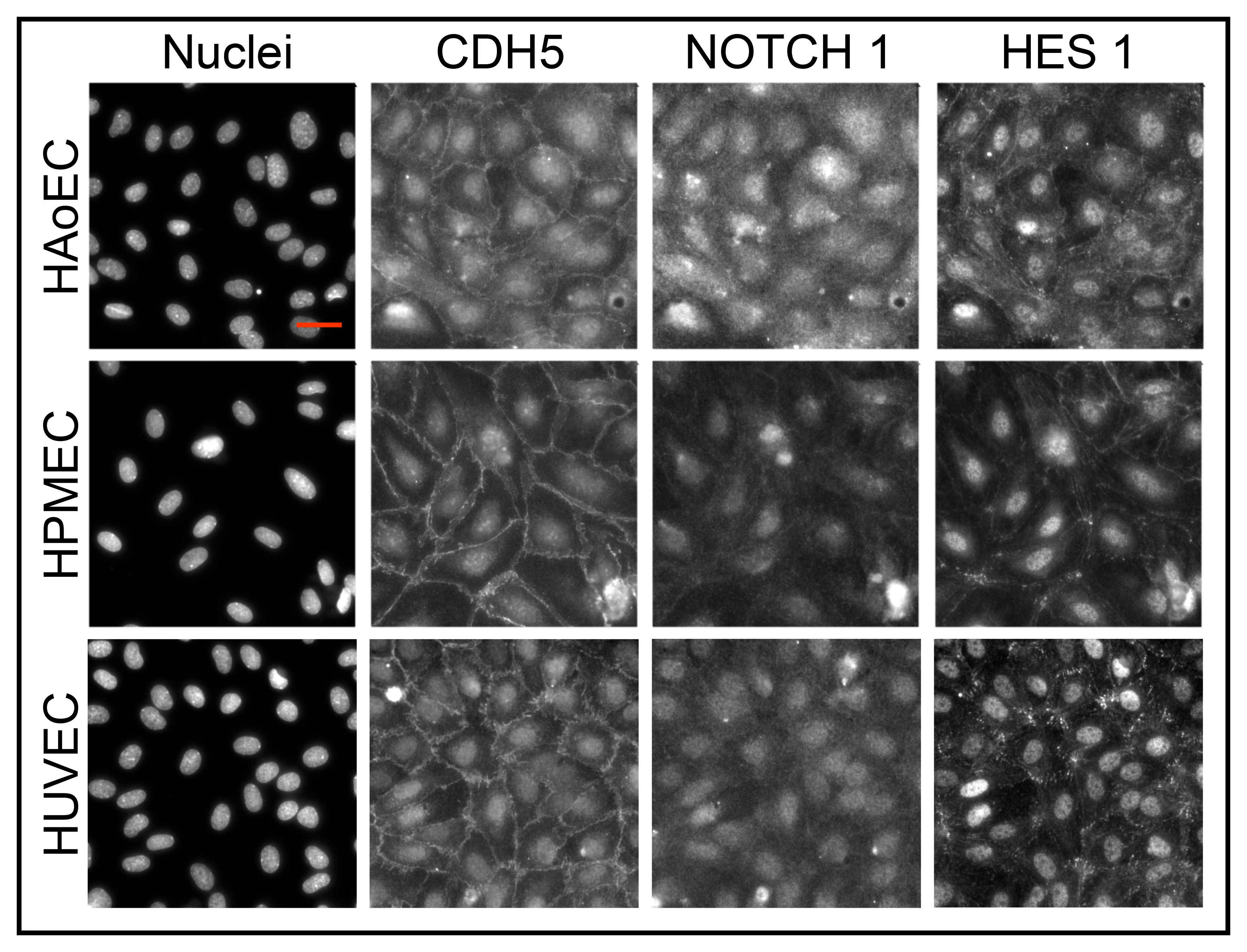
Overview of EC heterogeneity. Microphotographs comparing HUVECs, HAoECs and HPMECs stained for DNA (Hoechst, nuclei), VE-Cadherin (CDH5), Notch1 and Hes. Scalebar = 50 μm

EC junctions have been linked to specific function and phenotypes (Abu Taha et al., 2014; Lampugnani et al., 1995). Linear and stabilised junctions are present in mature quiescent phenotype; jagged and discontinuous junctions results from proliferation, migration or immature/mesenchymal phenotype; reticular patterns, observed in more stabilised junctions, can be also observed as transient structures (Kim and Cooper, 2018) and are associated with immune cell transmigration (Fernández-Martín et al., 2012).

In our study, HUVEC, HAoEC and HPMEC showed different junction patterns and cell morphology in standard *in vitro* culture conditions. Inspection of Notch1 and Hes1 staining intensity and localisation at the single-cell level appeared to be highly heterogeneous both intra-population and inter-population (Figure 1). Measuring and scoring individual EC phenotypes to demonstrate, quantify and explain EC heterogeneity at single cell resolution is a prohibitively tedious and biased approach if not automated. Therefore, we set out to create a dedicated platform that can seamlessly and comprehensively quantify variance of phenotypic parameters in cultured EC monolayers.

### Creation of a semi-automated image analysis pipeline for EC phenotyping

We developed a method to profile EC phenotypes at single cell level based on extracted high content analysis (HCA) features: Endothelial Cell Profiling Tool (ECPT, Figure 2 and Appendix 1). We used open-source software to generate an image analysis workflow able to take input from fixed or live images, with the aim of characterizing EC heterogeneity at the single-cell level (Fig. 2). Features and benchmark comparisons between ECPT and available tools is presented in Table 1. Fig. 2A demonstrates the unique capability of ECPT to precisely segment cells according to junctional CDH5 staining. Accurately segmenting cells is key to all downstream analyses and to evaluate cell junctions. However, CDH5 staining is very heterogeneous across different cell populations including those of different origins (e.g., HAoEC, HPMEC, HUVEC) and treatment (e.g., untreated vs VEGF treated EC). In terms of image analysis this renders standard thresholding techniques insufficient to appropriately contrast large collections of images for segmentation. To overcome this problem, we have developed a workflow in Fiji/ImageJ leveraging the Weka segmentation, a powerful machine learning (ML) tool. Fig. 2Ai is the original CDH5 image, Fig. 2A-ii are the corresponding output images following application of a Weka model based on an annotated dataset of 70 CDH5 images chosen randomly across our database of >20000 images. Fig 2 A-iii shows an overlay of individual junction objects. Individual junctions between two cells are precisely identified via ECPT and junction features are measured. Different junction classes were obtained by applying ML aided object classification (Cell Profiler Analyst, CPA using Fast Gentle Boosting) based on a junction’s measurements and colour code in Fig 2A-iii. Nuclei segmentation and downstream measurements were performed using standard Cell Profiler (CP) modules and thresholding methods (Otsu, Fig. 2B-i) to evaluate nuclear morphology, cell cycle (Figs. 2B-ii) and NOTCH signalling (Figs. 2C).

**Table 1:** ECPT features and comparison with other available software.

| Feature | Feature type | Software | ECPT_V1 | ECPT_V2 | Description | Benchmark |
| --- | --- | --- | --- | --- | --- | --- |
| PE Harmony/Columbus importer | Pre-processing | Fiji/ImageJ | YES | Improved | Imports images from PE Harmony and Columbus platform allowing custom labelling. Autodetects n of columns, rows, images, and channels. Allows performing image pre-processing (e.g., scaling, illumination correction). Improved speed in V2 | In native IA environment several of the following features are not available or limited. Processing could be done via python scripting to improve performance. |
| WEKA elaboration | Pre-processing | Fiji/ImageJ | YES | NC | Applies trained WEKA models to batch of images. Allow to select channel to be elaborated, model and to label class names for easier post-processing. | Arguably one of the best open-source pixel level segmentator using a familiar GUI (to ImageJ users) |
| WEKA models for CDH5 segmentation | Pre-processing | Fiji/ImageJ | YES | Improved | A WEKA model for junctions' segmentation trained on our full dataset. Efficiently segments junctions in all our images passing quality check but a new model might be required on different datasets. | Junction Mapper. Individual images need to be thresholded individually |
| Automatic image quality check | Pre-processing | Cell Profiler | NO | YES | In V2 we introduced an image quality check prior to object segmentation to avoid the need to manually select good quality images sets. | Same implementation as in Caceido Nat. Meth 2017 |
| Cell/nuclei/Junctions segmentation | OBJs detection | Cell Profiler | YES | Improved | CP modules to identify object of interest using standard CP modules. Segmentation is aided by intensity maps obtained through WEKA pre-processing. Improved junctions' identification in V2 due to improved WEKA model and restructuring of CP elaboration. V2 identifies Junctions as individual objects. | Standard OBJ segmentation similar efficiency in all platforms tested |
| Junctions Classification | OBJs class | CP analyst | YES | Improved | Classification of junctions according to measurements of junction objects. Improved in speed, accuracy, and quality of data by detection of whole junctional objects. | Not easily available in other software tested. CPA is user friendly, and parameters can be selected in a flexible way. Requires some experience with CPA for optimal results. |
| Cell classification | OBJs class | CP analyst | YES | NC | Classification of cells according to presence or absence of stress fibres | Not easily available in other software tested. CPA is user friendly, and parameters can be selected in a flexible way. Requires some experience with CPA for optimal results. |
| Extraction of spatial relationships information | OBJs detection | Cell Profiler | NO | YES | In V2 modules for establishment of neighbour relationship between individual junctions and cell object were introduced. This allows describing interaction between cells in terms of weighted interaction matrices essential for SAA. | Not available in any other tool to the best of author's knowledge |
| Detection of individual NOTCH puncta | OBJs detection | Cell Profiler | YES | NO | In V1 the measurement of individual NICD puncta was retained to mirror analyses in Wiseman et al. 2019. After restructuring of experimental and analysis setup we chose to remove this feature in favour of more robust nuclear intensity. | As for CDH5 based segmentation, WEKA allows achieving excellent results for pixel level segmentation. |
| Cell Maps export | Cross platform I/O | CP/Fiji/R | NO | YES | In V2 we introduced the possibility to share segmented objects across platforms which is key to exploit the potential of adespacial R package, i.e., to perform SAA across ~3000 images. | Not available in any other tool to the best of author's knowledge |
| R Elaboration | Data analysis | R Studio | YES | Improved | Scripts for all data processing performed in the work are provided as example allowing to reproduce all the data presented. Improvements in V2 include more consistent layout and scripts for SAA and relative visualisation. | Data I/O and elaboration in the R environment is more flexible than other platforms tested. E.g., SAA is impossible to implement is out-of-the-box tools tested by authors so far |

**Figure 2:**
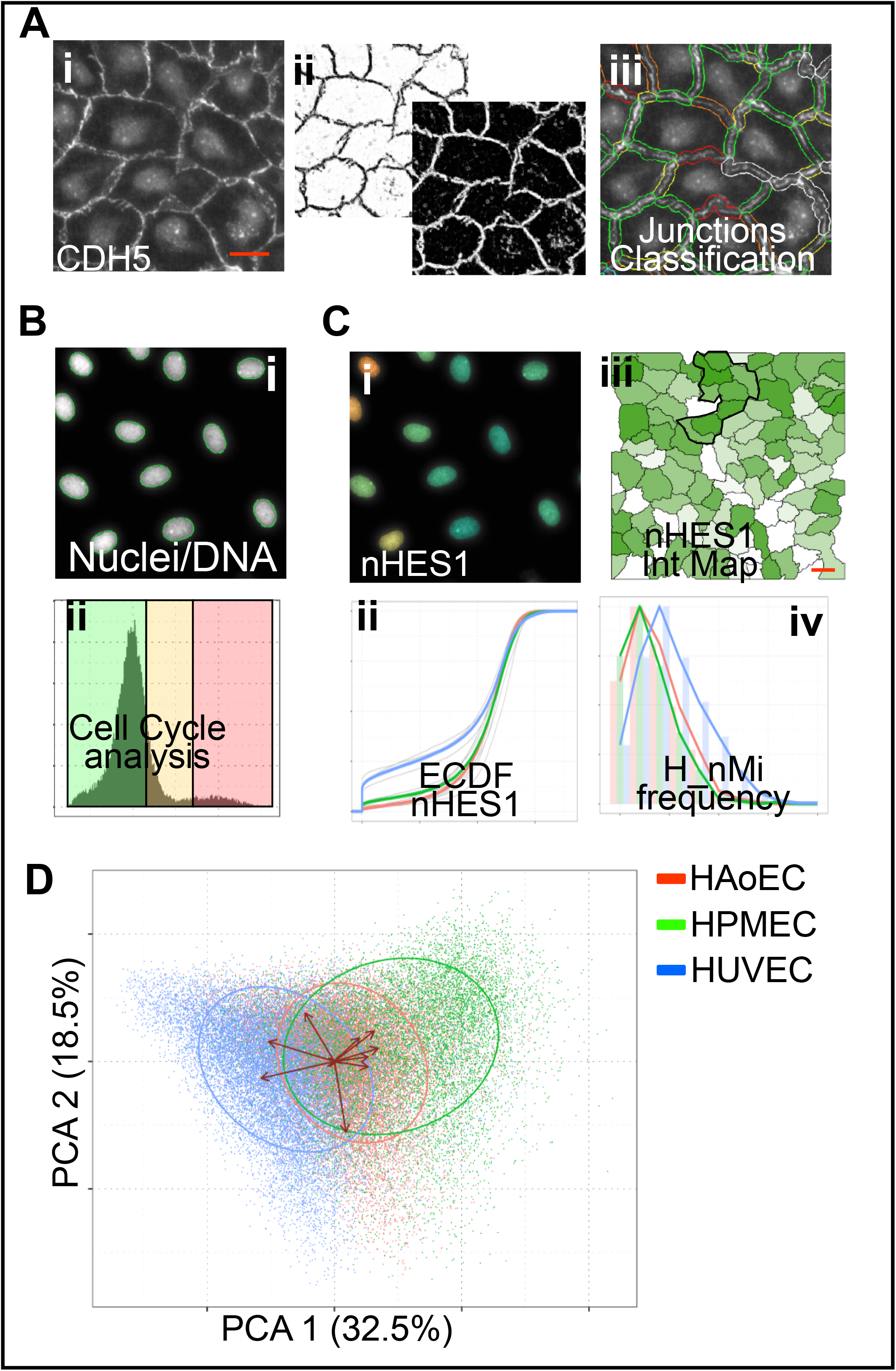
Image segmentation and ECPT features. **A**) Acquisition of high resolution (40X OM) images on each channel. CDH5 channel (i) is used to segment the cells and outline the cell junctions using Weka segmentator Fiji(ii), allowing individual junction and cells segmentation (iii). (**B**) Nuclei are stained with Hoechst (i) and the intensity is used to segment nuclei and measure DNA intensity to perform cell cycle analysis (ii). (**C**) Measurements of Hes1 nuclear signal intensity (i). Empirical cumulative distribution functions (ECDF) across all images by cell type (ii). Example nHES1 intensity map (iii) and frequency of cells with negative nLMi relative to HES1. **D)**Single cells PCA analysis on untreated EC cultured for 96 hours. Arrows indicate loading of different variables. Colours in C) and D) represent EC type. Scalebars = 50 μm

Overall, extending our previous proof-of-principle study (Wiseman et al., 2019) we chose features describing cell proliferation, morphology, spatial organisation, Notch activation and ML-aided classification of junctions. We then applied the ECPT workflow to a dataset composed of >20000 images from nine independent experiments obtaining a bulk of >300000 single cells across different cell types (i.e., HAoEC, HPMEC, and HUVEC) and experimental conditions (i.e., Initial cell density, time in culture, VEGF treatment).

We first asked whether HAoEC, HPMEC and HUVEC could be distinguished solely using the chosen features. We set to capture variance within our multivariate datasets, to identify discrete populations among heterogeneous cells and to understand which parameters had greater weight in defining cell phenotype. Principal components analysis (PCA) based dimensionality reduction cumulatively captured 51.1% of the variance (Figure 2D). HAoEC, HPMEC and HUVEC did not form distinct clusters and the cell populations were partially overlapping. Nevertheless, cell populations segregated according to the first two PCA components. This indicates that our parameters of choice capture key differences in the three EC populations. To understand the origin of this heterogeneity we dissected individual sets of features dependent on the same biological mechanism. We developed a dedicated user-friendly, open-source interface using the shiny package within Rstudio (https://shiny.rstudio.com) inspired by guidelines described previously (Lord et al., 2020). This shiny application allows subsetting through interactive and iterative selection of single cells or groups of cells for further comparison and analysis within R studio (Appendix 1 available online). We welcome readers to further explore our dataset available at https://github.com/exr98/HCA-uncovers-metastable-phenotypes-in-hEC-monolayers. To understand certain phenotypic differences in detail, we quantified variance of a few selected traits: proliferation, junction stability and NOTCH activation.

### Single cell Analysis of cell cycle

The integrated Hoechst nuclei signal intensity, commonly used as a proxy for DNA content in flow cytometry, was analysed based on previously published protocol (Roukos et al., 2015) to evaluate cell proliferation. The different stages of cell cycle (G1, S, G2/M) are thus defined via thresholding, with distinct peaks for G1 and G2/M (Fig. 3A). Late mitotic, (LM) cells, were detected by ML-aided object classification using features of Hoechst (DNA) and CDH5 stainings, including texture features (Fig. 3A). This automated method allowed to overcome specific limitations of previous approaches (Jones et al., 2009; Roukos et al., 2015).

**Figure 3:**
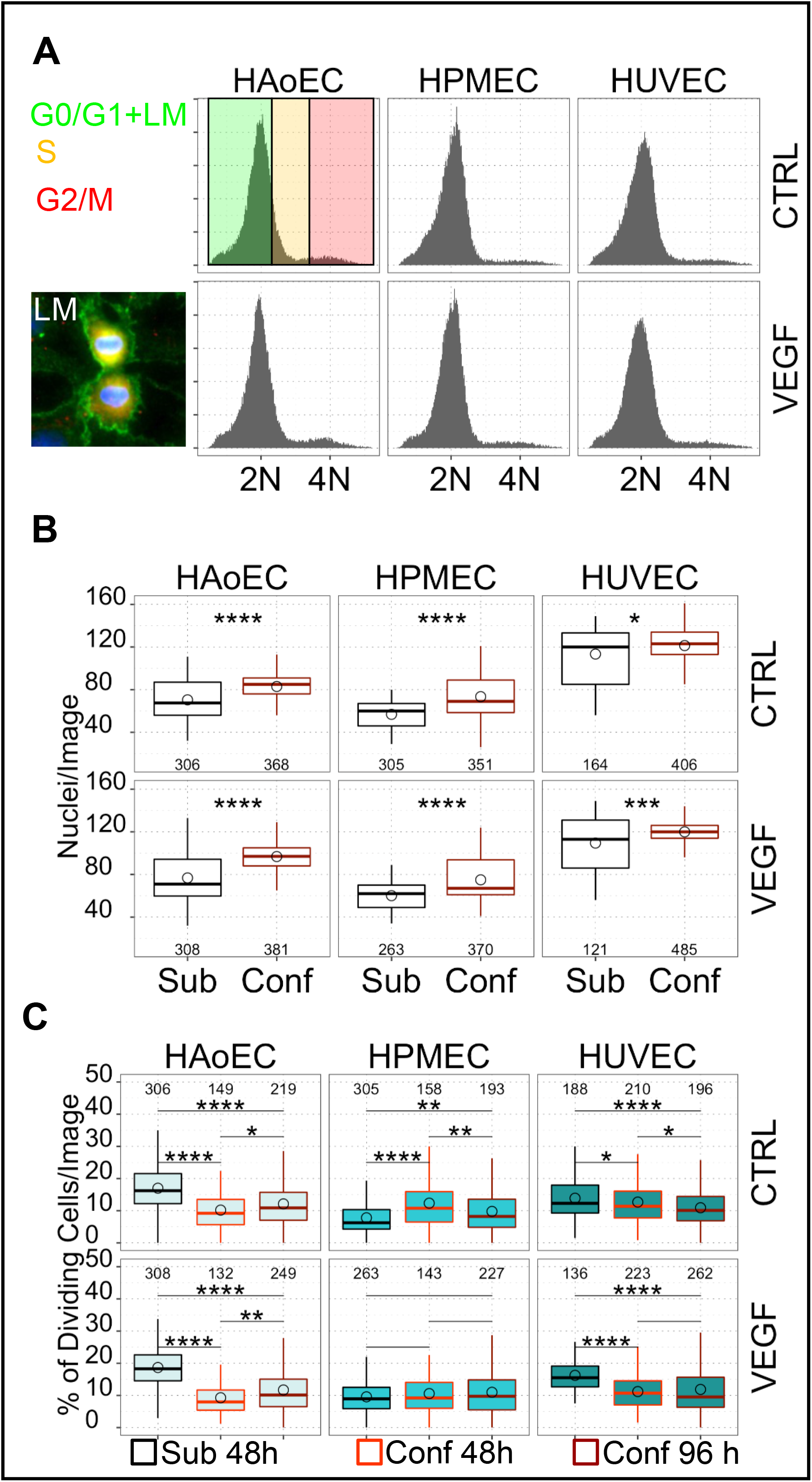
Cell cycle analysis. **A**) Histogram plots of DNA intensities by cell type (HUVEC, HAoEC, HPMEC) and treatments (CTRL, VEGF). Cells are classified in cell cycle phases depending on the signal intensity. Inset displays LM cells engaged in mitosis which were detected and quantified using CPA. **B**) Number of nuclei per image (cell density) in sub-confluent (black) or confluent (red) experiments for the 3 different EC types (HUVEC, HAoEC, HPMEC) and treatments (CTRL, VEGF) **C**) Percent of dividing cells (S, G2/M, LM) per image in HAoEC, HPMEC and HUVEC with or without treatment with 50 ng/ml VEGF in subconfluent or confluent conditions. n of observation for each statistical comparisons indicated as annotations in individual plots. * p<0.05, ** p<0.01, *** p<0.001, **** p<0.0001

Cell density is a well-established determinant of EC proliferation (Andriopoulou et al., 1999a; Bazzoni and Dejana, 2004) therefore we first set out to establish whether cell densities were homogeneous across cell types and different experiments.

By sub culturing HUVEC, HAoEC and HPMEC under the same standardised conditions, we noticed that these EC types present different intrinsic proliferation rates. We thus quantified proliferating cells and compared sub confluent and confluent cells which were also cultured for 48h or 96h upon seeding as longer time in culture has been shown to further stabilise the EC monolayer (Bazzoni and Dejana, 2004).

Proliferation was higher in sub confluent HAoEC and in HUVEC at baseline (average 16.9% and 13.8 % of dividing cells per field) and lower in HPMEC (average 7.8%). VEGF induced a small detectable increase in proliferation in all sub confluent EC (18.9%, 9.6% and 16.2% in HAoEC, HPMEC and HUVEC). In sub confluent cultures an average of 3.5% to 6.7% of cells were LM, demonstrating cells were effectively dividing at the time of the experiment.

In confluent cultures, the proliferation was lower in all EC except HPMEC which maintained values as in sub-confluent conditions. In confluent cultures percentage of cells in cell cycle ranged from 9.8% to 12.7% in all EC with little differences between CTRL and VEGF treated conditions. Longer culture conditions also had little effect on proliferation rates. In all confluent cultures examined the average percentage of LM cells dropped to values between 1.2% and 2%. Overall, our results are consistent with previous observations and demonstrate that EC proliferation is affected by cell density. Furthermore, confluent cultures under our conditions have small proliferation rates consistent with that observed *in vivo* (~4.8% of cells in S, G2 or M phases as measured by scRNA Seq of mouse aortic endothelium) (Lukowski et al., 2019)

Together these data validate the use of ECPT to characterise EC cell cycle under different experimental conditions. These are key quality controls when analysing cell phenotype and dynamic structures such as inter-endothelial cell junctions.

### Analysis of morphology and cytoskeleton arrangement

Cell morphology statistics were obtained directly from cell objects segmentation. Fig. 4A shows cell area comparisons across EC types and cell cycle stage in either sub-confluent (red, green, blue trace) or confluent (dashed grey traces) conditions. Overall, the three cell types had different average area and, as expected in sub confluent cultures, cell area increased with phases of cell cycle (between G0/G1 and G2/M phases) (Ginzberg et al., 2018). In confluent cultures cell area did not change across phases of cell cycle and confluent cells had lower proliferation rates. VEGF treatment did not induce significant differences in cell area in these conditions.

**Figure 4:**
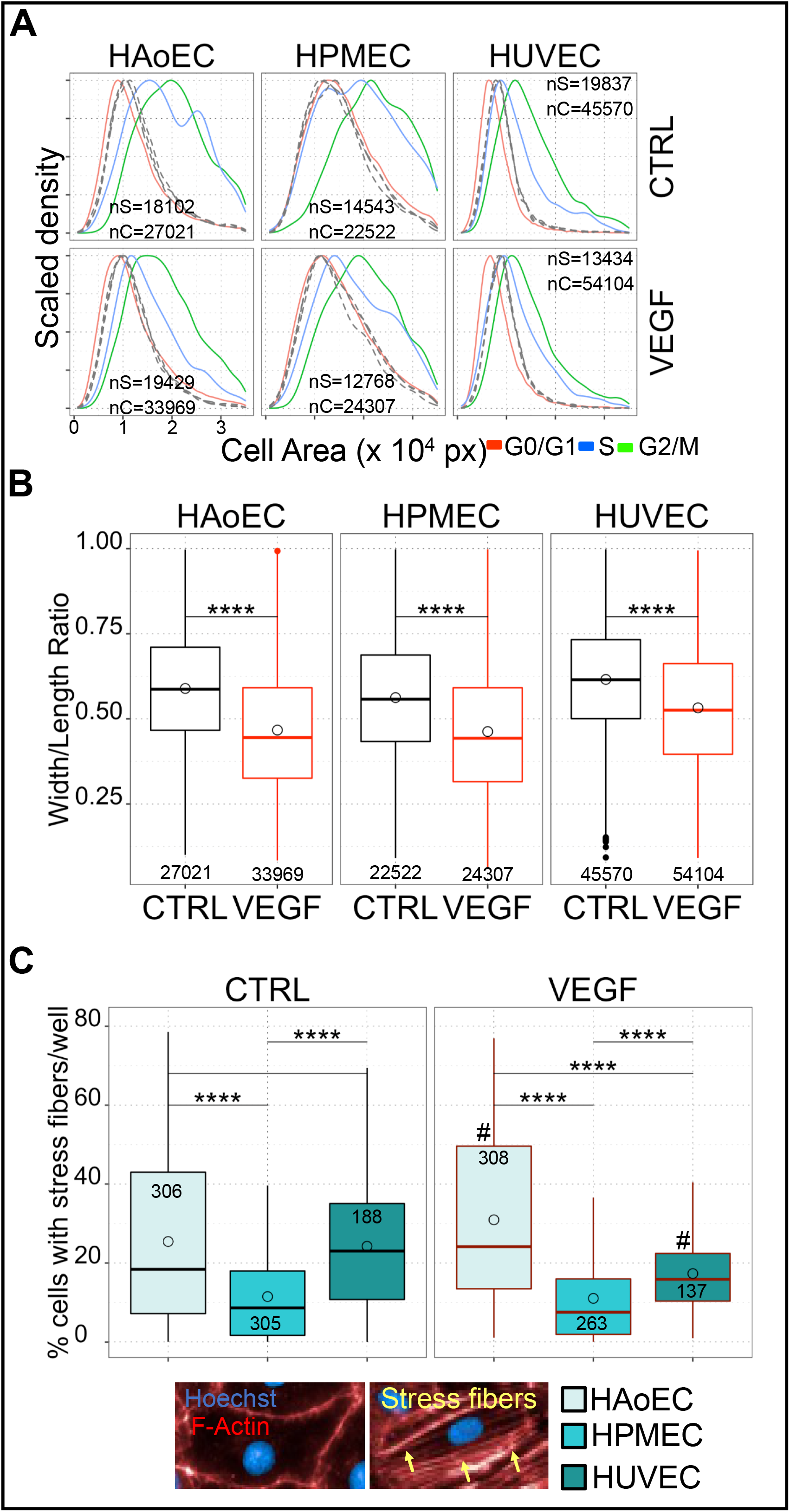
Analysis of cell morphology and cytoskeleton. **A**) Scaled density plots showing distribution of cell areas across cell types (HAoEC, HPMEC, HUVEC) and treatment (CTRL, VEGF) in sub-confluent (coloured traces, colour code as shown in legend) or confluent (dashed grey traces) conditions. **B)** Width to length ratio (WLR) of cell areas across cell types (HAoEC, HPMEC, HUVEC) and treatment (CTRL, VEGF). **C)** Analysis of stress fibres across cell types (HAoEC, HPMEC, HUVEC) and treatment (CTRL, VEGF). n of observation for each representation or statistical comparisons indicated as annotations in individual plots. * p<0.05, ** p<0.01, *** p<0.001, **** p<0.0001

Analysis of width to length ratio (WLR) in confluent cultures (Fig. 4B) shows a clear reduction in WLR in VEGF treated cells demonstrating cell elongation and confirming effectiveness of VEGF treatment. No difference was noted in WLR across different EC types or cell densities. By qualitative inspection of Phalloidin stained cells we noticed that different EC had qualitatively different distribution of actin cytoskeleton (see Abu Taha et al., 2014; Millán et al., 2010). In our sub confluent conditions HAoEC, HPMEC and HUVEC demonstrated different frequencies of cells with stress fibers (Fig. 4C). Data analysis showed that HAoEC and HUVEC had the highest proportion of cells with stress fibres while HPMEC had the lowest frequency. VEGF treatment induced a significant increase of these cell in HAoEC but a decrease in HUVEC, HPMEC maintained a low level like baseline.

Overall, the EC types analysed were remarkably different in terms of cell area and arrangement of cytoskeleton. As expected, cell area increased with progression of cell cycle in sub confluent cultures, but this effect was less prominent in confluent conditions where cells had lower yet appreciable proliferation rates. As expected, VEGF treatment clearly induced elongation in all cell types and induced cytoskeleton rearrangement in HAoEC and HUVEC.

### Analysis of junction heterogeneity

The description of EC junction dynamics and morphology has been of great interest due to the link with vascular permeability and angiogenesis. In general, non-proliferating EC are also those with continuous (“stabilised”) IEJ while migrating or proliferating EC demonstrate jagged or discontinuous junctions which are rapidly remodelling (Fernández-Martín et al., 2012; Millán et al., 2010). Average population measurements have been used by us (Veschini et al., 2011, 2007; Wiseman et al., 2019) and others to assess stability of EC monolayers which in turn correlate with functions such as trans-endothelial permeability to large proteins (Ferrero et al., 2004). Assessing IEJ at the single cell level enables to highlighting subtle nuances in inter-endothelial cells connectivity within the same monolayer. Nonetheless, measuring and classifying junctional signal by image analysis is technically challenging and no currently available software can evaluate multiple images in a standardised way.

To overcome this obstacle, we automated analysis of CDH5 pattern and junction morphology using the ML capabilities of CPA and integrated this into ECPT. Previous studies on EC junction (Brezovjakova et al., 2019; Seebach et al., 2015; Wiseman et al., 2019) did not analyse the proportion of different junction pattern; our method classifies whole junctions’ objects using an expert-trained ML algorithm (CPA with Fast Gentle Boosting, Appendix 1). In line with the literature in the field (Seebach et al., 2016), we classified junctions in a scale from 0 to 5 (Figure 5A). J0, J1 and J2 are discontinuous, highly jagged, or jagged junctions respectively. J3 and J4 are linear with J4 having continuous CDH5 signal distributed over a larger area than J3. J5 junctions are visually reminiscent of the reticular junctions previously described (Millán et al., 2010) but could also appear as transient structures (Kim and Cooper, 2018; Seebach et al., 2020). J0 junctions might result from mis-segmented cells therefore in downstream analyses we considered cells with less than 20% J0.

**Figure 5:**
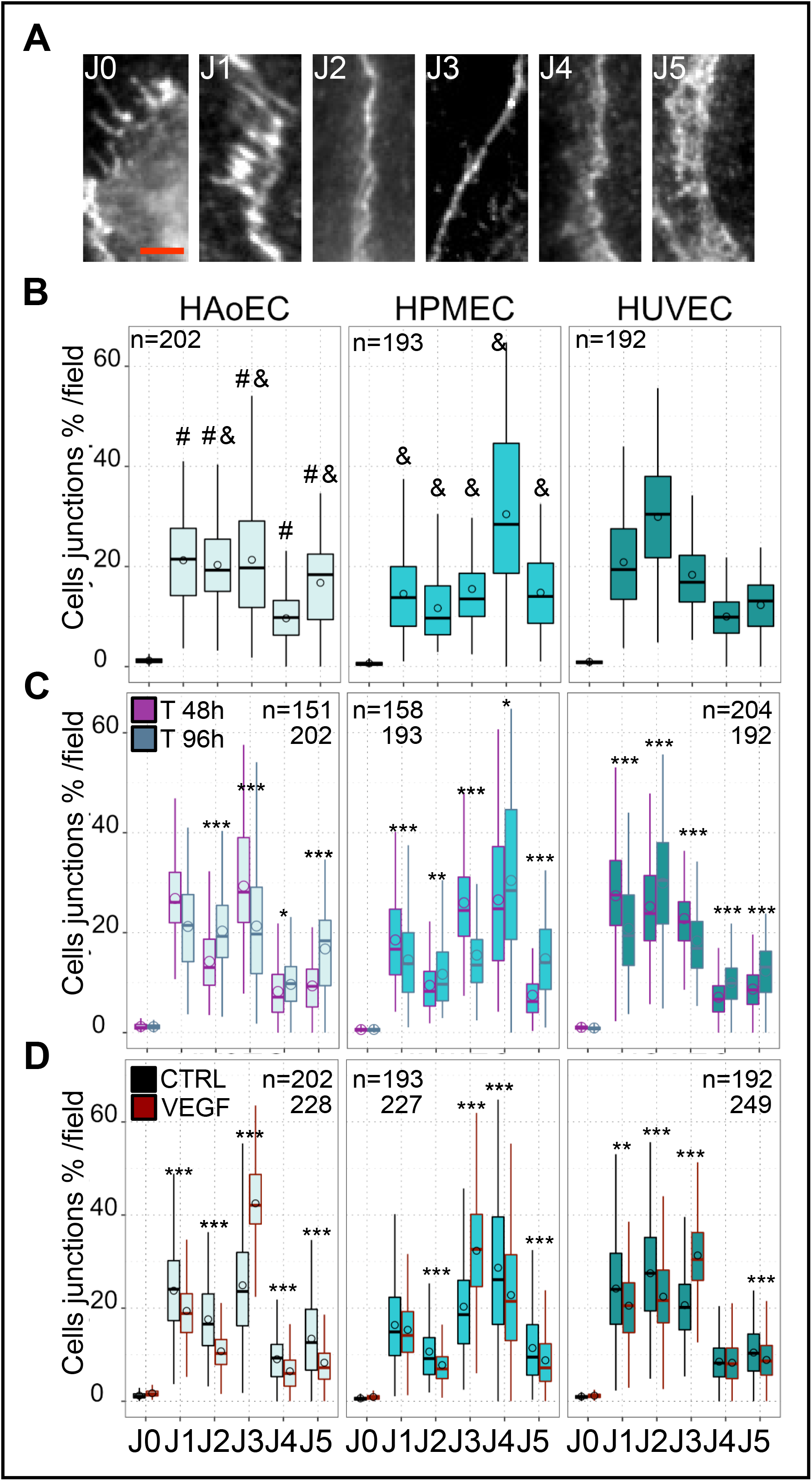
Image-based EC junction analysis. **A**) Examples of junctional structures imaged in EC monolayers. The junctions are classified as J0 (Not a junction or highly discontinuous), J1 (highly jagged and discontinuous), J2(Jagged), J3(Linear), J4 (linear reinforced signal) and J5 (linear/reticular) **B**) Percentage of cells in the different categories for across cell types under CTRL conditions and cultured for 96 hours **C**) Repartition of the cells in the different classes after 48h or 96h in culture. **D**) Percentage of cells in the different classes in basal conditions or after VEGF treatment. n of observation for each statistical comparisons indicated as annotations in individual plots. *** = p<0.001, & = p<0.001 against HPMEC, # = p<0.001 against HUVEC

Fig. 5B shows the average percent of junctions of each type across the three EC lines examined. HAoEC had a prevalence of J1, J2 and J3 types with scarce J4. HPMEC were characterised by a high proportion of J4 junctions while HUVEC had a high proportion of J2 type. Overall HPMEC and HUVEC had opposing phenotypes while HAoEC were more heterogeneous. Time in culture upon seeding have been shown to affect junction response to perturbation (Andriopoulou et al., 1999a), therefore we set to evaluate this effect under our experimental conditions. Fig. 5C shows the average percent of junctions of each type per cell per field comparing cells cultured for either 48h or 96h. The effect of longer culture in all cells was an increase in J4 and J5 types with corresponding decrease in J3. Furthermore, a decrease in highly jagged J1 with proportional increase in less jagged J2 type was also noted. Overall, we conclude that longer time in culture significantly and positively affects junction quality in line with previous reports (Andriopoulou et al., 1999b; Bazzoni and Dejana, 2004). Finally, we set to evaluate the effect of VEGF treatment in our system. As shown in Fig. 5D, the main effect of VEGF treatment on confluent cells cultured for 96h upon seeding was a sharp increase in J3 with correspondent decrease in all the other junction types.

Altogether these data demonstrate that the EC lines are very heterogeneous in respect to IEJ, and that many cells have a composition of different junctions which might also reflect differential connectivity with different neighbours. This junctional heterogeneity has been previously described and can now be detected and quantified by our ECPT. It suggests that IEJ architecture can be regulated locally as all cells were cultured under identical experimental conditions and all individual EC within monolayers exposed to the same environment. Overall, we showed that image analysis and supervised ML can be used to characterise EC junctions and highlight differences at the population level. Our workflow was able to efficiently distinguish changes in junction morphology after different time in culture or VEGF treatment and the junctional status of single cells can be used to study intra-population heterogeneity.

### Analysis of Notch activation

NOTCH signalling is a key modulator of EC development and function but assessment of signal heterogeneity in EC monolayers and its potential role in regulating these functions is currently lacking.

To assess heterogeneity in NOTCH signalling in our experiments we measured the intensities of intra-nuclear NOTCH1 (nN1) and intra-nuclear HES1 (nHES1) by ECPT. Fig. 6A illustrates the density distribution of the two signal intensities for all cells in our dataset at baseline demonstrating bulk differences between EC types. Analysis of mean signal intensity by field did not highlight striking differences across EC types and treatment. However, inspection of density distributions highlighted differences and suggests that these might originate in different repartitions of cells across signal intensities. To investigate this aspect, we binned signal intensities as illustrated by banding in Fig. 5A and calculated summary statistics by microscopic field for these bins. The results of these analyses for nN1 and nHES1 are shown in Fig. 6B and C. Analysis of nN1 highlighted differences in proportion of cells with either intermediate (2^8^) or high (2^10^) signal. At baseline HAoEC had the lowest percentage of cells in the intermediate intensity category and the highest percentage of cells in the high category. The opposite was true for HUVEC while HPMEC displayed similar percentage in the two categories and were intermediate between the other EC types. VEGF treatment set the moderate cells at intermediate levels for all cell types while it increased the percentage of high cells in HUVEC.

**Figure 6:**
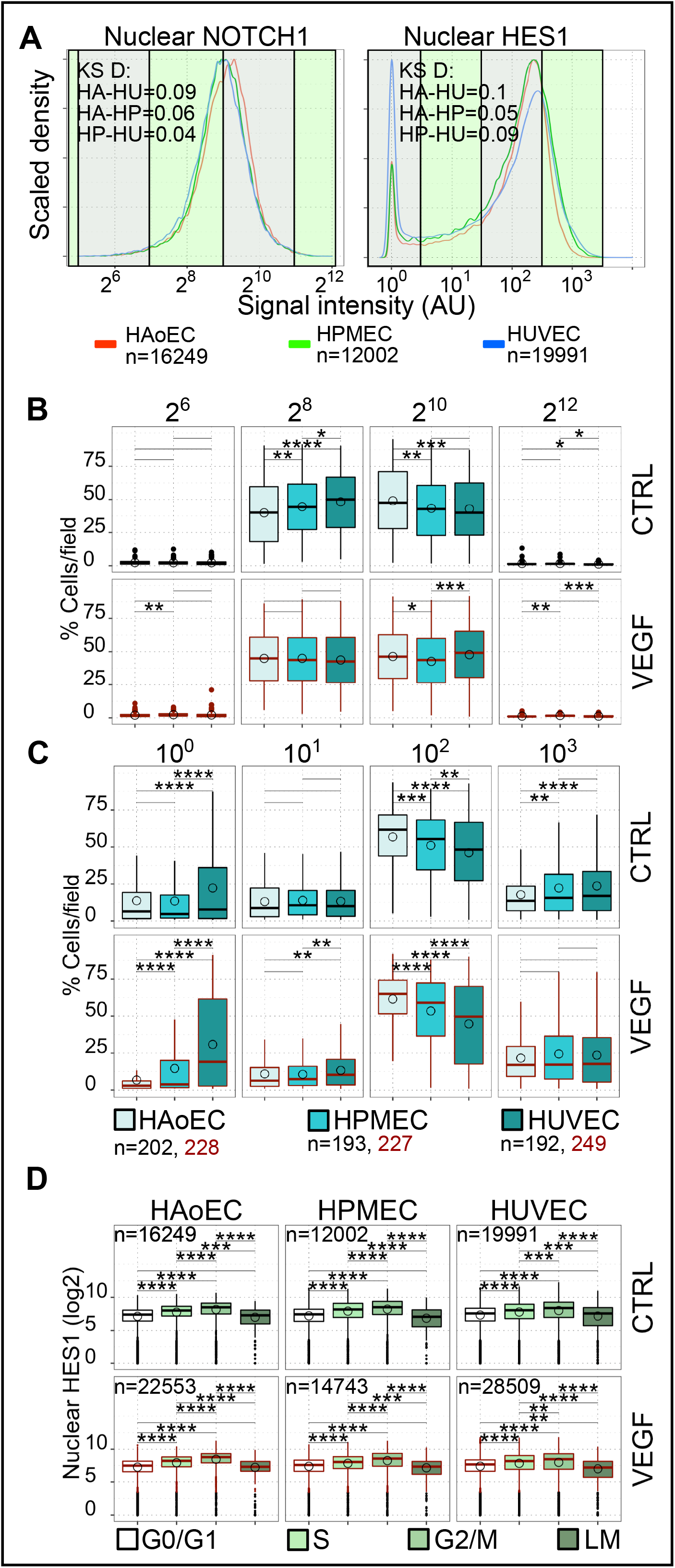
Analysis of NOTCH activation. **A**) Scaled density distributions of nN1 and nHES1 for HAoEC, HPMEC and HUVEC (coloured traces corresponding to legend). Distances between distributions are shown as annotations in plots (KS D, P<0.001 for all reported D). Green/transparent banding on plots indicates boundaries of selected intensity bins. **B)** Percent of cells per image (microscopic field) pertaining to different intensity bins (nN1) and EC types. **C)** Percent of cells per image (microscopic field) pertaining to different intensity bins (nHES1) and EC types. **D)** Intensity of nuclear HES signal (single cells) across phases of cell cycle, cell types (HAoEC, HPMEC, HUVEC) and treatment (CTRL, VEGF). n of observation for each statistical comparisons indicated as annotations under or in individual plots. * p<0.05, ** p<0.01, *** p<0.001, **** p<0.0001

Fig. 6C shows the analysis for nHES1 where cells were binned according to a log_10_ scale (bins referred to as low, intermediate, high, very high hereafter). HAoEC and HPMEC had both lower average percentage of cells in the low intensity category and greater percentage of cells in the high intensity category (13.7%, 56.9% and 13.5%, 51% in HAoEC and HPMEC respectively) in comparison to HUVEC (22.3% and 46.1%). VEGF treatment induced marked decrease in average percent of low intensity cells and corresponding increase in high intensity cells in HAoEC (6.8%, 61.5%) and had the opposite effect in HUVEC (30.8% and 44.7%).

We then evaluated the effect of differential time in culture by comparing nHES1 distribution in cells cultured for 48h or 96h. Suppl. Fig. 1A displays density distributions of nHES1 intensity which demonstrated marked changes in HAoEC and HUVEC and lesser differences in HPMEC. Suppl. Fig. 1B. shows that the overall effect of longer culture was a marked reduction of cells in the low intensity bin and a corresponding increase in the high bin in HAoEC and HUVEC while HPMEC were mostly unaffected.

Altogether, these results are in line with baseline gene expression levels of HES1 in the three EC type (Suppl. Fig. 2) and with previous gene expression studies (Chi et al., 2003) where arterial EC demonstrated higher levels of NOTCH signalling than venous EC. We also demonstrated previously unappreciated high levels of NOTCH signalling in HPMEC which were higher than HUVEC as also confirmed by gene expression analysis (Suppl. Fig. 2).

Importantly, our analysis highlights that 1) NOTCH signalling is not homogeneous in EC within monolayers, 2) all monolayers examined have similar ranges of nN1 and nHES1 signal intensities and, 3) bulk differences (e.g., qRT-PCR Suppl. Fig 2) result from different proportion of low, intermediate, or high signalling cells.

To estimate functional consequences of these observations we evaluated correlation between downstream NOTCH signalling and cell cycle. We found that the single cell levels of nHES1 were increased in cells engaged in cell cycle (~2-fold increase between G0/G1 and S and a further ~2-fold increase between S and G2/M) and decreased in dividing cells (~2 fold among LM and G0/G1) Fig. 6D. These observations were invariant across cell types and treatment and time in culture. It might be expected that cycling EC would have low NOTCH signalling as it has been previously shown that NOTCH signalling causes cell cycle arrest and inhibition of proliferation (Fang et al., 2017; Luo et al., 2021). In fact, all cell types in all phases of cell cycle had a significant proportion of cells with low nHES1 (Fig. 6D, shown as outlier dots in boxplots) and mitotic LM cells had an average/low level of nHES1.

We conclude from all data on nHES1 that, as expected and reported before (Fang et al., 2017), confluent EC have relatively high basal levels of NOTCH signalling. However, we also observed that within the same population some cells have low levels of nHES1. To interpret previous reports and our current data we hypothesize that sustained NOTCH signalling in confluent EC monolayers acts as a molecular break to limit cell cycle progression. Further, we postulate that it exists an escape mechanism by which cells can have low local levels of nHES1 allowing cell cycle finalisation despite sustained signalling at population level. Many previous studies have highlighted the role of NOTCH in mediating tissue patterning by mechanisms of lateral inhibition and in certain context lateral induction. We therefore set out to estimate the involvement of these processes using our ECPT.

### Spatial autocorrelation analysis of NOTCH signalling

To evaluate the role of lateral inhibition and lateral induction mechanisms we used spatial autocorrelation analysis (Moran’s analysis) to assess the distribution of nN1 and nHES1 signals in cell neighbourhoods and across entire monolayers. We first run the population level analysis (global Moran’s, as detailed in Suppl. Fig. 3 and methods) across all images in our dataset using either nN1 or nHES1 signals as inputs and length of junction object as the weighting parameter since estimated strength of neighbours’ interaction depends on extent of cell-cell contact.

Fig 7A shows that on average 25.7 – 40.4% of microscopic fields across cell types and VEGF treatment had statistically significant positive Moran’s Index (pGMi). Furthermore, 1.9 – 4.7% of microscopic fields had statistically significant negative Moran’s Index (nGMi) while 57.1 – 71.7% had randomly distributed cell intensities for nHES1. HUVEC had the highest percentage of microscopic fields with pGMI.

**Figure 7:**
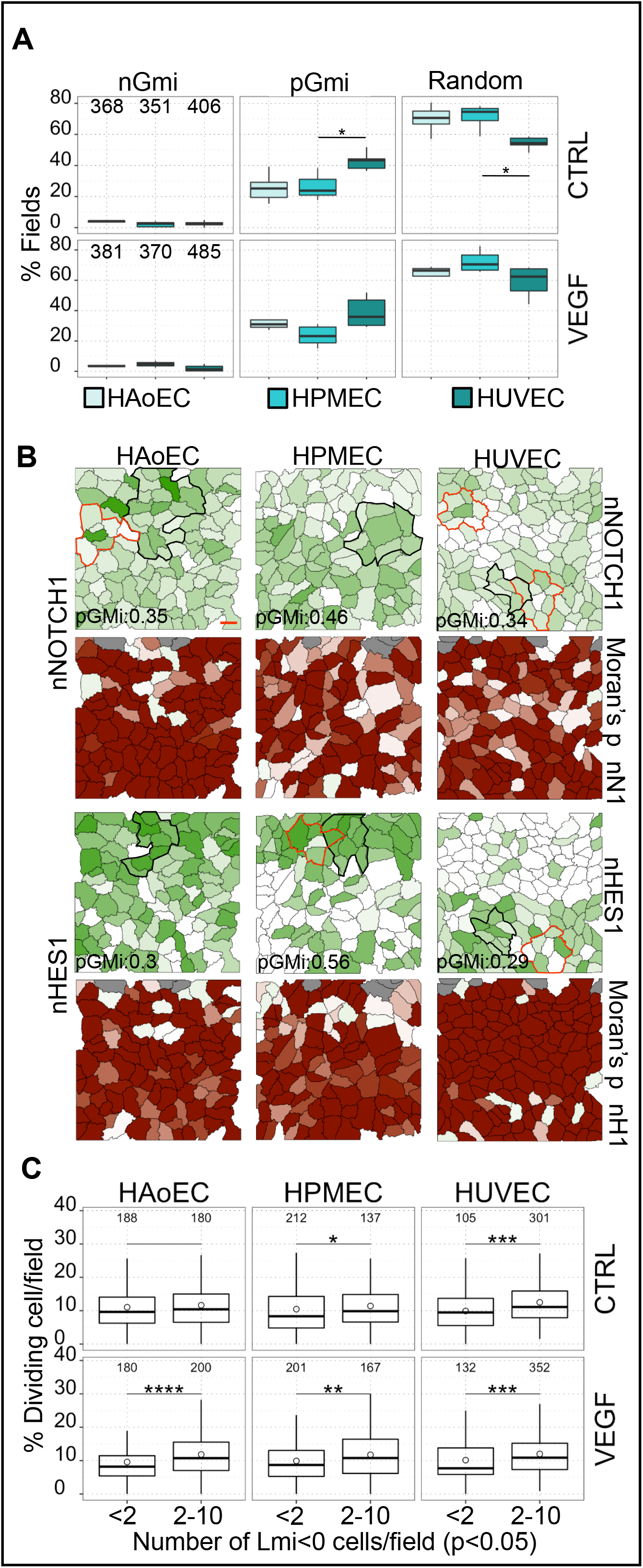
Spatial autocorrelation analysis. **A**) Box plots representing percent of fields with either significant (p<0.05) positive or negative global Moran’s Index (pGMi, nGMi) or random cell distributions (p>0.05 in both pGMI and nGMi) by cell type and treatment. **B**) Representative maps of cells with significant pMGi and relative Local Moran’s analysis. Green coloured maps refer to intensity of either nuclear NOTCH1 (log_10_) or HES1 (log_2_) signal (white=low intensity, dark green high intensity). Red coloured maps refer to p value of local Moran’s index LMi (white p<0.05, dark red p=1). Outlines in intensity maps indicated neighbourhood of selected cells with significant LMi associated with either positive (black) or negative (red) LMi. **C**) Boxplot (by EC type) representing percentages of dividing cells per images grouped in bins with low <2 or higher (2-10) numbers of cells with negative and significant LMi per field. n of observation for each statistical comparisons indicated as annotations under or in individual plots. * p<0.05, ** p<0.01, *** p<0.001, **** p<0.0001

Thus, the first key finding upon global Moran’s analysis is that most microscopic fields analysed (irrespective of cell type and treatment) had randomly distributed cell intensities highlighting the presence of intermediate cell states at the local population level. However, a sizeable fraction of the populations analysed shown evidence of clustered or sparse cell distributions suggesting that lateral inhibition and lateral induction are effectively active in the system. The average positive and negative global Moran’s indexes for all images analysed was ~0.4 and ~−0.3. These are intermediate values (extremes are 1 and −1 as illustrated in Suppl. Fig 3) which suggest that clustering or sparse distributions were detected in cell neighbourhoods smaller than the microscopic field analysed. Overall, global Moran’s analysis confirms that the observed bulk population distributions of nHES1 (Fig. 6A) are reflective of local arrangement after considering spatial relationships and suggests that clustering or sparseness can emerge in the wider population. To confirm and extend these observations we performed local Moran’s analysis which considers the immediate cell neighbourhood.

Fig. 7B illustrates one example of such analysis for both NOTCH (upper panels) and HES1 (lower panels). The intensity maps associated with nN1 and nHES1 highlight local heterogeneity in signalling and the p value maps indicate which cells in the map have statistically significant local Moran’s index (LMi, either positive or negative).

Comparisons of intensity and p value maps highlight clusters of cells with either similar or dissimilar signal values which were predicted by global Moran’s analysis and confirms that these clusters are smaller than the microscopic field (5-20 cells). 84% of all images had at least one cell with statistically significant negative LMI (nHES1) and 50% of all images had at least one cell with statistically significant positive LMI (nHES1). The frequency of nLMi were 1.6±1.2, 1.4±1.3 and 2.8±1.8 nLMI cells/per field in HAoEC and HPMEC and HUVEC. Average frequencies of pLMi were 0.6±0.8, 0.6±1 and 1±1.4 cells/field for HAoEC, HPMEC and HUVEC

VEGF treatment didn’t induce remarkable changes in nLMi or pLMi frequencies.

These observations confirm that of the global analysis and show that the distribution of nHES1 within EC monolayers is largely random and independent of cell type or exposure to VEGF treatment. However, monolayers are interspersed with neighbourhoods of cells that are engaged in lateral induction or inhibition processes as exemplified in Fig. 7B.

If cell proliferation is correlated with these phenomena as hypothesized above, we expect that cells able to progress through the cell cycle would be laterally inhibited to express lower levels of HES1 and escape cell cycle arrest. If that is the case, we would also expect to find more cells progressing through the cell cycle in areas where lateral inhibition is prevalent. To test this hypothesis within the limits of our experimental setup we compared microscopic fields with below- or above-average number of nLMi cells. Fig 7C shows the results of this analysis, confirming that microscopic fields with above average numbers of nLMI cells also had higher numbers of cells in cell cycle (Fig. 7C). As expected, no difference was found when considering abundance of pLMi cells.

Overall, we conclude that local lateral inhibition mechanisms can inhibit HES1 expression in a few cells within an EC population and thus increase the probability for individual cells to progress through the cell cycle.

Finally, as we had observed that longer times in culture quantitatively affected the distribution of nHES1 intensities (Suppl. Fig. 1A, and B) we sought to evaluate qualitative effects in Lmi. Suppl. Fig. 1C shows comparisons between counts of images containing 0 to 8 cells with negative Lmi (nLMi) and demonstrated that longer time in culture reduced the proportion of images with higher number of nLMi cells (without change in pLMi cell counts). Overall, longer time in culture reduced the number of cells engaged in lateral inhibition without subverting the overall distributions of cells with scattered presence of either small homogeneous cell clusters or laterally inhibited cells.

## Discussion

EC exert an outstanding variety of specialised functions (Augustin and Koh, 2017; Rafii et al., 2016) that are reflected in EC phenotypical heterogeneity. EC diversity arises during development and is preserved through homeostasis. The activation of differential genetic programs interplays with microenvironmental factors (Adams and Alitalo, 2007; Chi et al., 2003). Interestingly heterogeneity within the same vascular bed where microenvironmental factors are likely only undergoing minor fluctuations has been observed *in vivo* (McCarron et al., 2019) and deemed crucial to understand variable functions within the same EC type.

These mechanisms require similar cells to preserve discrete, diverse and adaptable phenotypes and the time scale at which these changes happen can be smaller than that of gene expression (Chapman et al., 2016; Stepanova et al., 2021). Thus, to justify rapid phenotype switches it is necessary to involve inherently dynamic processes such as junctional plasticity. Direct evaluation of dynamic changes in cell connectivity still poses considerable technical hurdles and despite great advances in live-microscopy, comprehensive cell phenotyping of single cells at the population level still relies heavily on end-point experiments. Although observation of fixed cells cannot provide information on the exact dynamics of the system, extensive profiling of single cells can inform about cell dynamic behaviours and help to develop hypotheses which can be evaluated experimentally or with computational methods (Altschuler and Wu, 2010).

To gain a better understanding of EC heterogeneity, connectivity, and emerging dynamic behaviours (McCarron et al., 2019, 2017) we developed ECPT and characterised three EC lines under standard conditions or supra-physiological levels of VEGF (like those observed in cancer). In line with previous studies (Chi et al., 2003), our data demonstrate that human aortic, umbilical vein and pulmonary microvascular EC present distinct phenotypes (Figs. 1–7) which are maintained at steady state, independently from microenvironmental stimuli. HAoEC, HPMEC and HUVEC displayed different intrinsic proliferative potential and demonstrated diffuse ability to proliferate, in contrast to the idea that EC undergo terminal differentiation and lose proliferative capacity (Yoder, 2018). Our image-based analysis of cell cycle based on DNA intensity highlighted interesting differences between the different EC lines in sub confluent conditions, with HUVEC and HAoEC proliferating faster than HPMEC, and an overall small increase in the dividing cell number after VEGF treatment. In confluent conditions all cell types settled to a lower proliferation rate which was similar for all EC types. It is well established that cytoskeleton arrangement correlates with differential EC phenotypes in different contexts. EC exposed to flow display stress fibers aligned with direction of flow and migrating EC display similar stress fibers along the direction of migration. Analysis of EC cytoskeleton (Fig. 4) demonstrated distinctive profiles in line with proliferation statuses in HAoEC, HPMEC and HUVEC, with microvascular HPMEC having the least stress fibres. At the same time, all cells demonstrated intra-population heterogeneity with intermixing of cells with or without stress fibres in individual monolayers (Fig. 4). This suggests a high degree of cellular rearrangement as indicated by time-lapse microscopy experiments showing that live EC in monolayers constantly rearrange their shape and connections (Kim and Cooper, 2018).

Differential EC functions leads to variance in IEJ which in turn regulate traffic of molecules and solutes in and out of capillaries and inhibit coagulation by preventing exposure of the underlying sub-intimal layer in arteries and veins. Analysis of IEJ in three different EC populations highlighted a high degree of junctional heterogeneity. In line with the data discussed above and existing literature, we show that HAoEC (arterial) had more linear junctions than HUVEC (venous) which instead tended to form highly dynamic jagged junctions. Importantly, HPMEC had the most linear/stabilised junctions while HAoEC had more heterogeneous jagged/linear junctions and stress fibres (Fig. 4). This is reminiscent of the unique role the pulmonary and arterial vessels play in regulating solute exchange or adapting to high flow, respectively. This is in line with recent sc-RNA sequencing data demonstrating that >40% of murine HAoEC express mesenchymal genes (Lukowski et al., 2019) and are therefore expected to display morphological and junctional heterogeneity accompanied by stress fibres.

In agreement with results reported before (Bazzoni and Dejana, 2004) longer culture times and VEGF treatment in our experiments induced characteristic changes in cell morphology and promoted more continuous junction types across all EC types.

Distinctive features among different EC populations seem to be hard coded within cell gene expression program as cells from individual donors had vastly overlapping phenotypes and the features were maintained in all lines upon several passages in culture.

If the decision to enter cell cycle or to maintain stable junctions were dependent on clear-cut differential gene expression, we would expect to find a limited collection of homogeneous phenotypes. However, we find a continuous range of EC phenotypes within the same EC monolayer. To interpret EC intra-population heterogeneity, we focussed on NOTCH signalling for its well-established role as a coordinator of the opposing functions of EC proliferation (Fang et al., 2017; Luo et al., 2021) and junctional complex stabilisation (Bentley et al., 2014).

Furthermore, the NOTCH pathway is one of the main drivers of endothelial cell heterogeneity and is linked to vascular maturation, arteriovenous specification and angiogenic sprouting (Fish and Wythe, 2015; Hellström et al., 2007; Potente et al., 2011; Torres-Vázquez et al., 2003). Although NOTCH signalling dynamics have been studied in HUVEC and retina, it is still unclear how it affects EC heterogeneity in different organs. Nuclear NOTCH1 (N1-ICD) acts as a transcription co-factor to induce the expression of target genes such as the HES/HEY family of transcription factors and is therefore active in the nucleus. In line with previous results on HEY1/HEY2 expression in HUVEC (Aranguren et al., 2013), we hypothesised that EC within monolayers would have a homogeneous level of NOTCH signalling that was either low or absent in venous EC and high in arterial EC. One study has reported appreciable baseline NICD levels in HPMEC by western blotting (Zong et al., 2018) but not in comparison with other EC lines. Recent single-cell RNA sequencing data of human lungs has highlighted heterogeneity among arterial, venous, and microvascular EC, which displayed intermediate phenotypes (Kalucka et al., 2020). Interestingly, activation of NOTCH downstream target genes also seems heterogeneous across and within EC populations (Travaglini et al., 2020). Fluctuations in NOTCH signalling have been hypothesised in EC monolayers and demonstrated in EC during sprouting angiogenesis where NOTCH acts as a bistable switch (Ubezio et al., 2016). It has been demonstrated that leading tip cells induce NOTCH signalling in trailing stalk cells via Dll4-NOTCH1 leading to lateral inhibition of the tip cell phenotype (Ubezio et al., 2016) whereas EC within a stabilised monolayer cannot acquire a tip cell phenotype and all EC receive in principle similar NOTCH stimulation from neighbours.

By measuring NOTCH1 and HES1 in individual EC, we found that different EC within their monolayer have different levels of NOTCH activation which is also reflected in different bulk gene expression measures of NOTCH target genes (HES1). However, the clearest differences across cell types and treatments were found in relative proportions of cells with differential signal within the same EC population, suggesting that individual cells could acquire differential phenotypes which can be modulated by VEGF treatment (Fig 6). This degree of heterogeneity in NOTCH signalling in neighbouring cells is also strongly suggesting that NOTCH phenotype is regulated dynamically in EC monolayers with lateral inhibition and lateral induction being two potential candidate mechanisms. To investigate this hypothesis, we performed spatial autocorrelation analysis (Fig. 7) which demonstrated high degree of heterogeneity of NOTCH and HES1 in EC within the same monolayer If either lateral inhibition or lateral induction were the prevailing mechanism in regulating the NOTCH phenotype in EC, we would expect sparse, or uniform spatial distribution of NOTCH and HES1 activation. However, we found a high degree of randomness in the spatial distribution of these signals. Alternatively, the two mechanisms might act in concert to produce qualitatively different cell distributions which seems plausible as all EC types analysed expressed both Dll4 and Jagged1 ligands (Suppl. Fig. 2) which have been involved in lateral inhibition and lateral induction respectively. When we performed local spatial autocorrelation analysis we confirmed that both mechanisms seem to occur together and our results are in line with previous mathematical models (Boareto et al., 2016) showing that concurrent lateral inhibition and induction can generate intermediate cell states. Our current data demonstrate that intermediate phenotypes in both nN1 and nHES1 are common in all EC analysed, irrespective of treatment, and that lateral induction or lateral inhibition patterns emerge locally. This complex spatial distribution does not seem compatible with stabilised phenotypes and rather suggests a scenario where cell phenotype is dynamically regulated and can change over time to exert differential functions while maintaining a stable balance across the wider population.

Novel spatialised mathematical models of NOTCH signalling could be used in future work to assess whether NOTCH is sufficient to generate the heterogeneity we found in our experiments or if further layers of regulation need to be accounted for.

To evaluate the functional consequences of our findings we asked whether spatial patterning accompanied by differential signalling could affect other parameters in our dataset. We did not find any correlation between the extent or spatial distribution of nN1 and nHES and junction status, possibly because junctions are remodelled at a very fast pace in cultured EC (Kim and Cooper, 2018) and thus our experimental context using fixed cells might fail to resolve processes at this timescale. When we considered cell proliferation, we found that HES1 was, on average, higher in cells progressing through the cell cycle but several cells had low levels irrespective of their cell cycle status. When we compared the abundance of laterally inhibited cells with dividing cells at the population level, we consistently found that populations containing more of these cells were also proliferating at slightly higher pace. Overall, our data suggest that the decision to initiate and progress through cell cycle in continuous monolayers with high basal levels of NOTCH signalling is regulated by the extent (and possibly duration) of lateral inhibition in the local cell neighbourhood. Moreover, we show that the formation of patches of cells with similar NOTCH signalling is a common finding in EC monolayers *in vitro*. It will be interesting to investigate whether this is reflected *in vivo* and the functional consequences of this phenomenon considering previous work has identified patches of cells with differential Ca++ signalling and density of endothelial M3 muscarinic acetylcholine receptors (AchRM3s) *in vivo* (McCarron et al., 2019, 2017). However, *in vivo* investigations are still posing significative technical challenges to high throughput analyses such as those presented here.

It will be also interesting in future work to evaluate whether the patterns observed in our experiments are stable or remodelled over time and the timescales of these changes. The ECPT presented in this work is a step forward in comparison to available platforms (see Table 1) and represents a bridge between experimental and computational setups where experimental ECPT data can be used to guide development of spatialised mathematical models which in turn can be used to guide further experimentations. The experimental setup introduced in the present work does not extend to resolving highly dynamic processes, however ECPT paves the way towards quantifying more advanced experiments such as measurements of gene transcription by FISH or live imaging using fluorescent reporters.

Overall, the ECPT allows for the evaluation of cell-intrinsic mechanisms of monolayer maintenance and plasticity, excluding variability caused by microenvironmental factors, in line with previous observations on blood vessel heterogeneity *in vivo* originating from EC rather than perivascular cells (Chavkin and Hirschi, 2020). It will be important to elucidate in future studies how the crosstalk with perivascular cells and microenvironment can affect emerging endothelial behaviours.

We envisage that coupling our ECPT workflow with live imaging setups, computational modelling, and single cells transcriptomic analysis will open the way to a much deeper understanding of emerging dynamic endothelial behaviours and thus help to develop novel more effective therapies for regenerative medicine, prevention of cardiovascular diseases and the treatment of cancer.

## Materials and Methods

### Cell culture

HAoEC, HPMEC and HUVEC (PromoCell) were plated on 10 μg/mL fibronectin (from human plasma, Promocell)-coated flasks, grown in EGMV2 medium (Promocell) in the absence of antibiotics, detached with Accutase (Thermo Fisher Scientific, Waltham, MA), and used by passage 5. We analysed two distinct donors for each cell type which were chosen excluding diseases affecting the vasculature (e.g., diabetes). Donor’s age (HAoEC, HPMEC) was between 50 and 63 years. For experiments, 4 × 10^4^ EC per well were seeded in fibronectin-coated 96-well plates (μclear, Greiner) and cultured for 48 or 96h under basal (EGMV2, Promocell) or activated (EGMV2 + 50 ng/mL VEGFA, Peprotech, London, UK) conditions in triplicate paralleling conditions described previously (Andriopoulou et al., 1999a). The EC formed confluent monolayers at microscopic inspection (phase contrast, 10x-20x OM) at the time of immunostaining and image acquisition.

### Immunostaining

Cells were fixed with 2% paraformaldehyde in phosphate buffered saline (PBS) for 10 minutes at room temperature. Cells were blocked 1h with PBS supplemented with 1% fetal bovine serum (FBS) and permeabilised with 0.1% Triton X 100. Cells were then incubated for 1h at room temperature with primary antibodies against CDH5 (Ve-Cadherin Novusbio NB600-1409, 1 μg/mL final), NOTCH1 (Abcam, ab194122, Alexa 647-conjugated, 1 μg/mL final) and Hes1 (Abcam, ab119776, 1 μg/mL final). Plates were washed and incubated 1h with 1 μg/mL secondary Alexa 488-conjugated and Alexa-555-conjugated antibody (Thermo), Hoechst 33342 (1 μg/mL, Sigma) and Phalloidin-Atto 647N (Sigma).

### Image Acquisition

We obtained images from slides with an Operetta CLS system (PerkinElmer, Waltham, MA) equipped with a 40×water-immersion lens (NA 1.1). In each well, 3 areas were acquired. Each area is composed of nine microscopic fields at 40× magnification (Supplementary Figure 1). We standardised acquisition parameters (led power, exposure time) throughout different experiments and used HUVEC as a standard for calibration in all experiments. We analysed an image database containing 28000 images (7000 fields in four fluorescence channels) extracted from nine independent experiments conducted on EC lines from two different donor each. Two intraexperiment replicates were conducted for each experiment.

### Endothelial Cell Profiling Tool

We used a combination of machine learning-aided image segmentation (ImageJ) (Schindelin et al., 2012) and an image-based cell profiling tool (CellProfiler) (Carpenter et al., 2006) to extract the phenotype of single EC in monolayers. Our workflow enables to measure EC morphology (Area, perimeter, shape descriptors, cell neighbours), NOTCH1, HES1, CDH5 and DNA intensities and to characterise inter-endothelial junctions (IEJ). In the present study, we chose to analyse only selected features with recognised functions in EC biology. Image texture features were measured and only used during the training of machine learning algorithms (Caicedo et al., 2017; Jones et al., 2008) which in turn were used to classify junction morphology and LM cells. ECPT scripts and methods including FIJI/ImageJ macros for image importing and pre-processing are detailed in Appendix 1.

We Imported and cleaned the results into R studio excluding artifacts and mis-segmented cell objects (extreme values in cell Area or signal intensity, NAs in measurements). We then calculated continuous and categorical (Cell Cycle) parameters. Following guidelines suggested by Caceido et al. (Caicedo et al., 2017) we pre-processed the database to exclude miss-segmented cells and to normalise the measurements prior to dimensionality reduction or single factor analysis. We reformatted and tidied the database and calculated summary statistics using packages from the Tidyverse packages collection (Dplyr, Tidyr, Tibble, Forcats, Purr, Ggplot2) We conducted PCA analysis on ECPT parameters (Cell morphology, intensity, neighbourhood and junctional status, Supplementary Table 1) using the prcomp function (R Stats package) generating 12 PCA components. All plots in figures are generated using the Ggplot2 R package.

We created a shiny application (https://CRAN.R-project.org/package=shiny) for interactive selection and visualisation of data of interest. All files containing the code required to reproduce al the plots in the paper and to run the shiny application using our dataset are available on https://github.com/exr98/HCA-uncovers-metastable-phenotypes-in-hEC-monolayers.

### Western blotting

Cells were scraped in the presence of RIPA buffer (Sigma-Aldrich) containing protease (Millipore, UK) and phosphatase inhibitors (Sigma-Aldrich, UK), left on ice for 15 min, and centrifuged for 5 min in a refrigerated microfuge. Supernatants were assessed for total protein using the BCA protein quantitation kit (Thermo Fisher Scientific, UK). 15 μg of protein were separated on NuPAGE 4-12% Bis-Tris gels (Invitrogen) before being transferred to nitrocellulose membranes (GE Healthcare, Amersham, UK). After probing with primary and secondary antibodies, membranes were developed using Clarity™ Western ECL Substrate (Biorad, UK) and read using a ChemiDoc system (Biorad). Antibodies for Notch 1 (Abcam, ab194122), Hes1 (Abcam, ab119776), VE-Cadherin (Novus bio NB600-1409), and β-tubulin (Cell Signaling Technology, UK). Goat anti-mouse and anti-rabbit horseradish peroxidase (HRP)-conjugated antibodies were from Dako (Agilent).

### RNA extraction and qRT-PCR (quantitative reverse-transcription Polymerase Chain Reaction)

Total RNA was extracted and purified using the Monarch total RNA miniprep kit according to the manufacturer’s instructions. The resulting RNA was quantified using Nanodrop (ISOGEN Life Science). For real-time PCR, 1 μg of RNA was used for reverse transcription using the iScript cDNA synthesis kit (Bio-rad). The gene expression analysis was carried out using the SsoAdvancedTM Universal SYBR Green Supermix (Bio-rad) and analyzed by means of a Stratagene Mx3000P (Agilent Technologies) in real time, primers used are listed in Table 2 (Supplementary material).

### Statistical analysis

To compare multiple groups, we used one-way ANOVA followed by Tukey’s HSD post-hoc test. We considered p< 0.05 (*) statistically significant and p<0.01 (**), p<0.001 (***) and p<0.0001(****) highly significant.

To evaluate linear correlation between continuous variables we calculated Pearson’s Correlation coefficient I and considered p<0.001 as highly statistically significant.

Comparisons between intensity distributions were performed by two-sided Kolmogorov Smirnov (KS) test as implemented in the “stats” R package. We considered comparisons with p<0.01 to be statistically significant and reported corresponding distance index (D).

Statistical significance in global and local Moran’s analyses is computed by random permutations as implemented in the adespatial R package we used 999 permutations in each test. We considered p<0.05 in Gmi or Lmi to be statistically significant (Dray, 2011).

## Acknowledgments

We wish to thank Dr Eng. VCG La Ferla for creating the FIJI importer macro including GUI.

## Data availability

All raw and elaborated data are available at https://github.com/exr98/HCA-uncovers-metastable-phenotypes-in-hEC-monolayers. To facilitate data exploration and re-use we have developed a dedicated data browser using the Shiny app environment. Original images are available upon request to the authors.

---

**Supplementary Figure 1: Effect of long culture conditions on NOTCH activation. A**) Scaled density distributions of nN1 and nHES1 for HAoEC, HPMEC and HUVEC (coloured traces corresponding to 48h or 96h culture). **B)** Percent of cells per image (microscopic field) pertaining to different intensity bins (nN1) and EC types. **D)** Counts of fields according to numbers of nLMi cells/field, cell types (HAoEC, HPMEC, HUVEC) and treatment (CTRL, VEGF). * p<0.05, ** p<0.01, *** p<0.001, **** p<0.0001

**Supplementary Figure 2: Gene expression analysis:** Analysis of expression of selected genes in HUVEC, HPMEC and HUVEC. Two donors for each cell type, measurements for donors are assembled. CDH5 (pan-endothelial marker), LYVE1 and PROX1 (lymphatic EC markers), DLL4, NOTCH1, JAG1 (NOTCH signalling), HES1 (NOTCH target gene). n=6 for all cell types for all genes except LYVE1 and PROX1 (n=8). * p<0.05, ** p<0.01, *** p<0.001, **** p<0.0001

**Supplementary Figure 3: Vignette of Moran’s analysis. A)** Cell maps for each individual field are extracted by ECPT. Cell neighbours and length of neighbour-neighbour junctions (x,y,z) are recorded into a spatially weighted matrix (SWM) of dimensions nxn (n is the number of cells in the respective field). Test parameter values (x’, y’, z’, w’, e.g., nH1 or nHES1 intensities) are recorded into a vector of length n. For each cell local Moran’s values are computed by evaluating variance in signal with neighbours weighted by extent of interaction. **B)** Prototypic examples of global Moran’s analysis in regular cells distributions. Random distribution of intensities yields pGMi and nGMi close to 0 (i). Sparse distributions yield pGMi close to 0 and nGMi approaching 1 (ii). Clustered distributions yield pGMi approaching 1 and nGmi close to 0 (iii).

**Supplementary Figure 4: Validation of antibodies specificity. A)** Western Blot (WB) using ab119776 antibody against HES1 overlayed with tubulin, B) WB using NB600-1409 antibody against CDH5 overlayed with tubulin. C) WB using ab194122 against NOTCH1 intracellular domain. WB resolves a band 98kDa as expected from cleaved NOTCH1, whole NOTCH1 protein is not resolved by WB under our experimental conditions. All WB are performed using lysates from HUVEC, HAoEC and HPMEC as indicated.

