## Supplementary Figures and Tables for "High content Image Analysis to study phenotypic heterogeneity in endothelial cell monolayers"

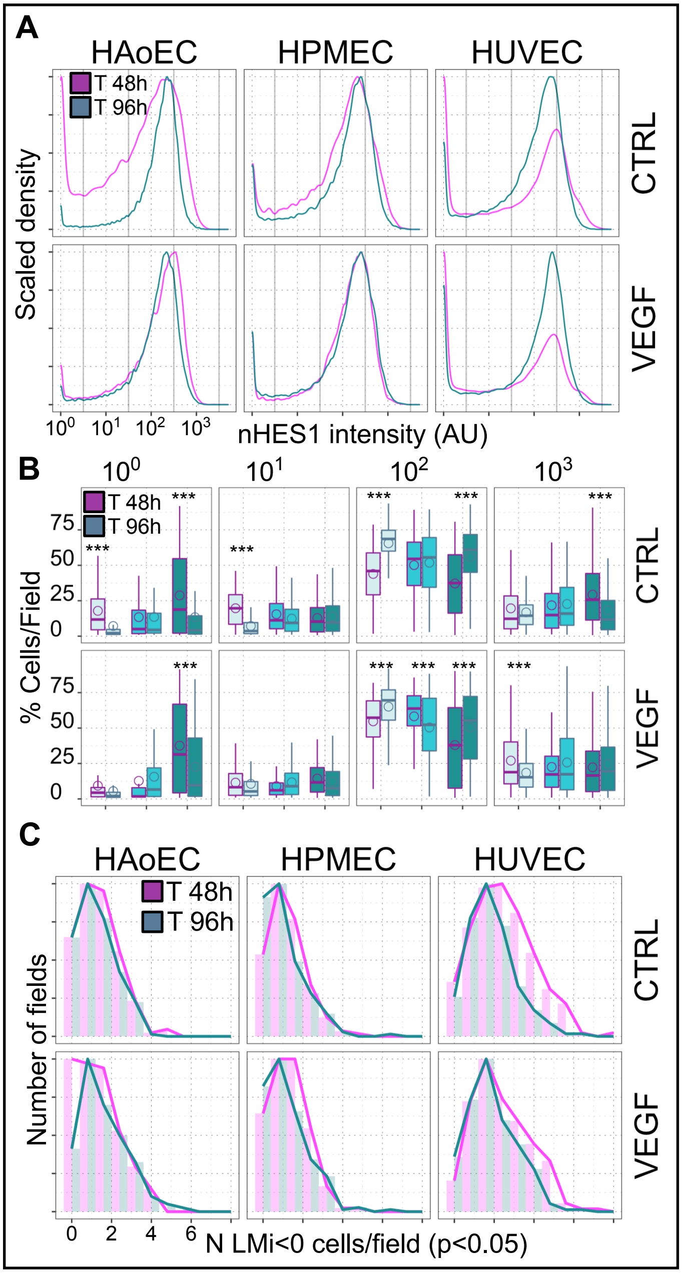


**Supplementary Figure 1: Effect of long culture conditions on NOTCH activation.**

**A**) Scaled density distributions of nN1 and nHES1 for HAoEC, HPMEC and HUVEC (coloured traces corresponding to 48h or 96h culture). **B)** Percent of cells per image (microscopic field) pertaining to different intensity bins (nN1) and EC types. **D)** Counts of fields according to numbers of nLMi cells/field, cell types (HAoEC, HPMEC, HUVEC) and treatment (CTRL, VEGF). * $p<0.05$, ** $p<0.01$, *** $p<0.001$, **** $p<0.0001$

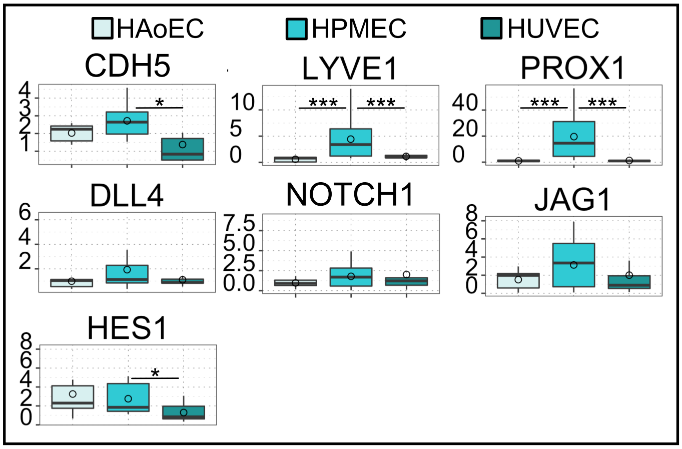


**Supplementary Figure 2: Gene expression analysis:** Analysis of expression of selected genes in HUVEC, HPMEC and HUVEC. Two donors for each cell type, measurements for donors are assembled. CDH5 (pan-endothelial marker), LYVE1 and PROX1 (lymphatic EC markers), DLL4, NOTCH1, JAG1 (NOTCH signalling), HES1 (NOTCH target gene). n=6 for all cell types for all genes except LYVE1 and PROX1 (n=8). * p<0.05, ** p<0.01, *** p<0.001, **** p<0.0001

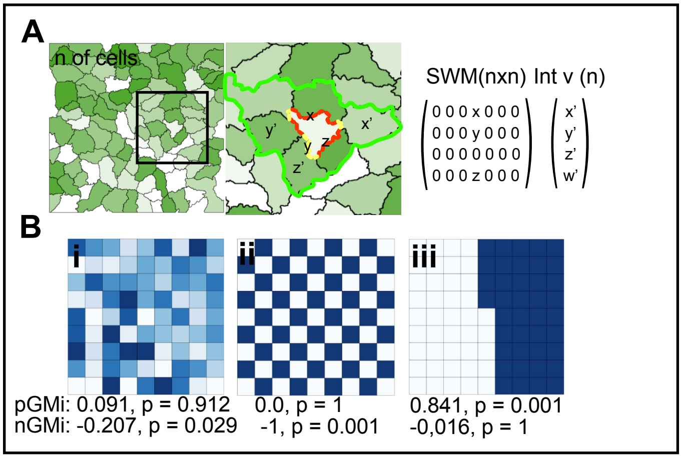


**Supplementary Figure 3: Vignette of Moran's analysis. A)** Cell maps for each individual field are extracted by ECPT. Cell neighbours and length of neighbour-neighbour junctions (x,y,z) are recorded into a spatially weighted matrix (SWM) of dimensions nxn (n is the number of cells in the respective field). Test parameter values (x', y', z', w', e.g., nH1 or nHES1 intensities) are recorded into a vector of length n. For each cell local Moran's values are computed by evaluating variance in signal with neighbours weighted by extent of interaction. **B)** Prototypic examples of global Moran's analysis in regular cells distributions. Random distribution of intensities yields pGMi and nGMi close to 0 (i). Sparse distributions yield pGMi close to 0 and nGMi approaching 1 (ii). Clustered distributions yield pGMi approaching 1 and nGmi close to 0 (iii).

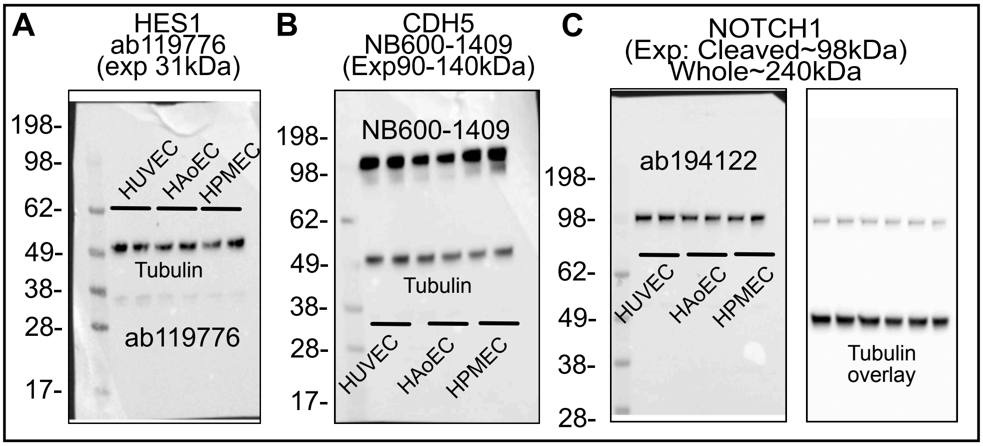


**Supplementary Figure 4: Validation of antibodies specificity. A)** Western Blot (WB) using ab119776 antibody against HES1 overlayed with tubulin, B) WB using NB600-1409 antibody against CDH5 overlayed with tubulin. C) WB using ab194122 against NOTCH1 intracellular domain. WB resolves a band 98kDa as expected from cleaved NOTCH1, whole NOTCH1 protein is not resolved by WB under our experimental conditions. All WB are performed using lysates from HUVEC, HAoEC and HPMEC as indicated.

| Primer | Forward sequence | Reverse sequence |
|---|---|---|
| RPL19 | CAGAAGATACCGTGAATCTAAG | TGTTTTTGAACACATTCCCC |
| CDH5 | CGCAATAGACAAGGACATAAC | TATCGTGATTATCCGTGAGG |
| LYVE1 | AGGCTCTTTGCGTGCAGAA | GGTTCGCCTTTTTGCTCACAA |
| PROX1 | AAAGGACGGTAGGGACAGCAT | CCTTGGGGATTCATGGCACTAA |
| DLL4 | GTCTCCACGCCGGTATTGG | CAGGTGAAATTGAAGGGCAGT |
| NOTCH1 | GAGGCGTGGCAGACTATGC | CTTGTACTCCGTCAGCGTGA |
| JAG1 | GTCCATGCAGAACGTGAACG | GCGGGACTGATACTCCTTGA |
| HES1 | TCAACACGACACCGGATAAAC | GCCGCGAGCTATCTTTCTTCA |

**Table 2: Primer sequences used in qRT-PCR.**
